# Freely accessible ready to use global infrastructure for SARS-CoV-2 monitoring

**DOI:** 10.1101/2021.03.25.437046

**Authors:** Wolfgang Maier, Simon Bray, Marius van den Beek, Dave Bouvier, Nathaniel Coraor, Milad Miladi, Babita Singh, Jordi Rambla De Argila, Dannon Baker, Nathan Roach, Simon Gladman, Frederik Coppens, Darren P Martin, Andrew Lonie, Björn Grüning, Sergei L. Kosakovsky Pond, Anton Nekrutenko

## Abstract

The COVID-19 pandemic is the first global health crisis to occur in the age of big genomic data.Although data generation capacity is well established and sufficiently standardized, analytical capacity is not. To establish analytical capacity it is necessary to pull together global computational resources and deliver the best open source tools and analysis workflows within a ready to use, universally accessible resource. Such a resource should not be controlled by a single research group, institution, or country. Instead it should be maintained by a community of users and developers who ensure that the system remains operational and populated with current tools. A community is also essential for facilitating the types of discourse needed to establish best analytical practices. Bringing together public computational research infrastructure from the USA, Europe, and Australia, we developed a distributed data analysis platform that accomplishes these goals. It is immediately accessible to anyone in the world and is designed for the analysis of rapidly growing collections of deep sequencing datasets. We demonstrate its utility by detecting allelic variants in high-quality existing SARS-CoV-2 sequencing datasets and by continuous reanalysis of COG-UK data. All workflows, data, and documentation is available at https://covid19.galaxyproject.org.

## Introduction

Effectively monitoring global infectious disease crises, such as the COVID-19 pandemic, requires capacity to generate and analyze large volumes of sequencing data in near real time. These data have proven essential for monitoring the emergence and spread of new variants, and for understanding the evolutionary dynamics of the virus.

Two sequencing platforms in combination with several established library preparation strategies are predominantly used to generate SARS-CoV-2 sequence data. These sequencing data are therefore mostly harmonized and available in defined formats from standard repositories, e.g. NCBI SRA in the US and ENA in Europe. However, data alone do not equal knowledge: they need to be analyzed. Here we focus on the foundational task of identifying fixed differences and minor allelic variants in viral genomes from deep sequencing data (resequencing).

The current analytical landscape for SARS-CoV-2 genomic data is not well harmonized. A review of published literature shows that different research groups perform sequence analyses in distinct ways with some approaches being more appropriate than others (see Ref.1 for examples). The discrepancy between the state of data generation and data analysis–a defined set of experimental strategies diverging into a multitude of analytical approaches–makes it hard to integrate newly acquired knowledge and compare results across studies.

A number of well-designed and validated SARS-CoV-2 data analysis approaches already exist. For example, the ARTIC network^2,3^ provides best practices for the analysis of amplicon resequencing datasets. There comprise precisely documented analyses and validated open source software for identification of sequence variants, phylogenetic reconstruction, analysis of selection, and so on. However, software tools form just one component of the analysis ecosystem. The sheer size of already existing and continuously generated new data requires adequate computational infrastructure to utilize these software tools. Access to appropriate infrastructure is problematic for many research groups even in developed countries, as it requires expertise in resource procurement, configuration, and maintenance. The existence of commercial cloud computing resources does not fully address this situation, because these resources still need to be configured and funded. In addition, computational clouds are predominantly owned by US-based companies and many countries have policies that make paying foreign cloud providers difficult. The challenges are even more acute outside of industrialized Western countries, where robust research computing infrastructure simply does not exist and researchers have no means to pay for computation.

Fortunately, in the United States, the European Union, and Australia powerful public computational infrastructures have been designed specifically for research purposes on a global scale. These include the XSEDE4 consortium in the US, the deNBI5 and ELIXIR6 consortia in the EU, and Nectar Cloud7 in Australia. Large scale public computing resources are ideally suited for tackling the informatics challenges of the current pandemic because they are globally accessible, support diverse configuration schemes (from traditional computational clusters to fully virtualized cloud-like setups), and provide the full spectrum of cutting edge hardware. However, because these resources have traditionally been used primarily for physical and engineering sciences, they remain under-utilized by biomedical researchers.

This public computational infrastructure coupled with open-source software tools offer a complete solution to SARS-CoV-2 data analytics challenges. All that is presently missing is *glue* to bind these into a unified analysis platform capable of fluidly managing users, allocating storage, and pairing analysis tools with appropriate computational resources. Furthermore such a platform would need to accommodate researchers with a spectrum of computational expertise by providing both a graphical user interface as well as programmatic access. Finally, such a platform should *not* be developed by a single PI, group, or an institution. Instead it should be supported by the international community of users, developers, and educators.

Here we used the global Galaxy platform^8,9^ to build such a resource. It is a public genomic data analysis portal geared to monitoring public infectious disease emergencies matching the scale of the COVID-19 pandemic. Galaxy combines computational infrastructure from the US, EU, and Australia to deliver a truly global, distributed, scalable, and free system. We demonstrate the utility of this system by analyzing large (thousands of samples) existing datasets derived from RNAseq and Ampliconic SARS-CoV-2 sequencing experiments to show that (1) large read-level datasets can be analyzed on public computational infrastructures in a matter of hours, and (2) outcomes of these analyses can be readily transformed into publishable results by incorporating current information about emerging SARS-CoV-2 variants and genome sites that are evolving under selection.

## Results and Discussion

### Open software for open science

Our system is housed on three globally distributed public Galaxy instances in the US (http://usegalaxy.org), the EU (http://usegalaxy.eu), and Australia (http://usegalaxy.org.au). Each is capable of supporting thousands of users running hundreds of thousands of analyses per month. Anyone can create an account and obtain immediate access to as much computation as one might reasonably need (with a limit on the number of concurrent analyses) and 250 Gb of disk space, which can be increased based on the needs of an individual user.

An important element of our platform is its integration of training modules. These are powered by the Galaxy Training Network^10^ (GTN; https://training.galaxyproject.org), and already include interactive SARS-CoV-2 analysis tutorials.

The system relies on three principal software systems: Galaxy, Jupyter, and ObservableHQ (Fig. 1) organized into two analysis stages. Stage 1 is the most computationally intensive, taking as input raw sequencing data and yielding intermediate summaries of these data such as lists of allelic-variants (hereafter called AVs to distinguish allelic-variants from SARS-CoV-2 genomic variants) and annotations (similar to VCF files). Stage 2 focuses on the interpretation and visualization of the Stage 1 outputs.

**Figure 1.**
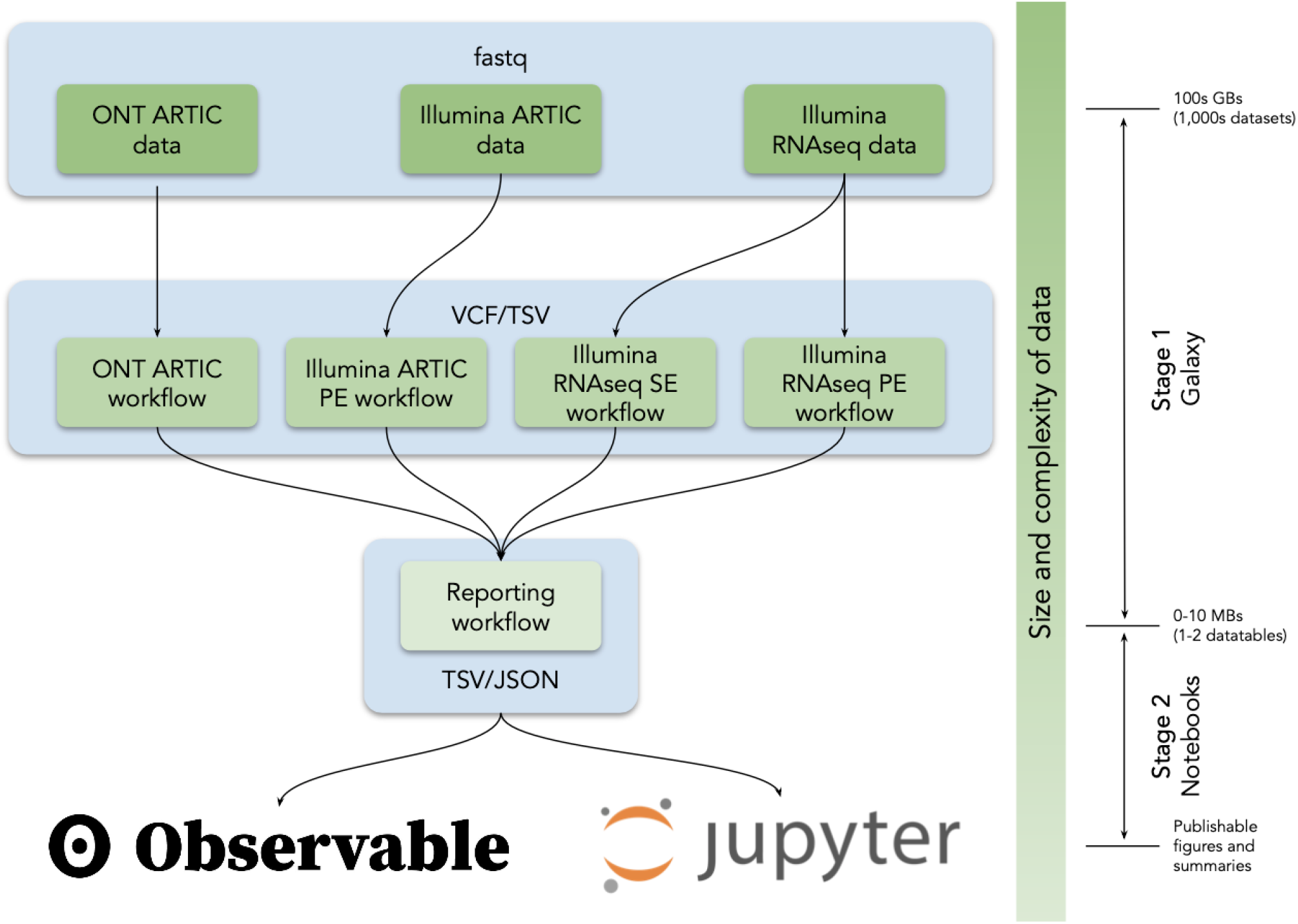
Analysis flow in our analysis system. VCF = variant call format, TSV = tab separated values, JSON = JavaScript Object Notation.

The software components of Stage 1 are all well defined, mature, and robust utilities within the Galaxy tool ecosystem, including open source utilities for sequencing quality control, read mapping, assembly, and AV calling. The Galaxy tool ecosystem hosts thousands of open source tools and is closely aligned with the BioConda project^11^ to facilitate versioning and distribution. Stage 1 analyses are performed entirely in Galaxy on public infrastructure.

The software components of Stage 2 are widely used analytics and visualization environments such as Jupyter, RStudio, or ObservableHQ. These environments allow users to explore data using a notebook paradigm where snippets of code (cells) are used for data transformation and visualization. While Jupyter and RStudio are heavily used in biomedical research, ObservableHQ is a new serverless application that relies on JavaScript for data processing directly in a browser. It is ideally suited for rapid development of rich interactive visualization dashboards.

Galaxy provides all tools, computational infrastructure, graphical interface functionality and programmatic access via a well-defined API for analyses performed in Stage 1, whereas for Stage 2, Galaxy directly integrates and supplies all necessary computational resources for user-invoked Jupyter and RStudio^12^ notebooks. Galaxy provides data formatted for ingestion by ObservableHQ notebooks and interacts with these via stable URLs that are dataset specific.

### A ready-to-use SARS-CoV-2 allelic-variant surveillance system

To demonstrate the utility of a free globally accessible analysis environment, we developed five analysis workflows to support the identification of SARS-CoV-2 AVs from deep sequencing reads (Table 1). A user begins the analysis by uploading reads in FASTQ format into Galaxy (Fig. 1) as a dataset collection (a dataset collection is a way to represent an arbitrarily large collection of samples as a singular entity within a user’s workspace; see Ref.^13^). These datasets can either be uploaded by the user, obtained from local data mirrors or retrieved directly from the Sequence Read Archive at NCBI. The four primary analysis workflows (#1-4 in Table 1) convert FASTQ data to annotated AVs through a series of steps that include quality control, trimming, mapping, deduplication, AV calling, and filtering. All Illumina workflows use `lofreq`^14^ as the principal AV caller. We selected it based on extensive testing (Methods). All four workflows produce identically annotated VCF output that is further processed by the Reporting workflow (#5 in Table 1) to generate data tables describing AVs. These data tables are further processed with Jupyter directly in Galaxy and with ObservableHQ to generate all figures and tables such as those shown in the paper (links are given in Table 2). All workflows, data, and “how-to” documentation is available at https://covid19.galaxyproject.org.

**Table 1.**
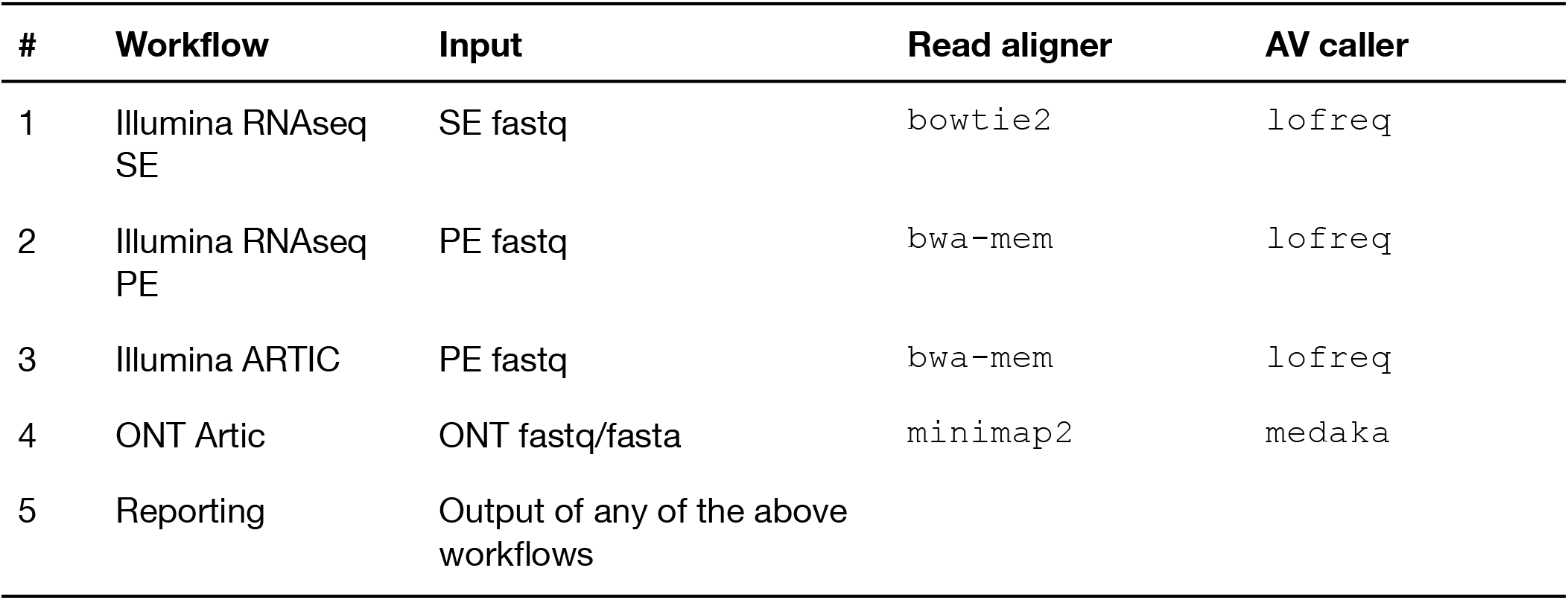
Description of analysis workflows. PE = paired-end; SE = single end

**Table 2.** Jupyter and ObservableHQ notebooks using for Stage 2 of the analysis

| Notebook framework | Figure from this paper generated using this notebook | Link |
| --- | --- | --- |
| Jupyter | 4, S1- S6 | GitHub \| Colaboratory |
| ObservableHQ | 3 (also see Table S2) | ObservableHQ |

### Application to characterizing intrahost allelic-variants in existing data

As of late January 2021 the NCBI sequence read archive contained 190,288 raw read SARS-CoV-2 datasets (Fig. 2). There are three primary types of data: (1) Illumina-based Ampliconic, (2) Oxford nanopore (ONT)-based Ampliconic, and (3) Illumina-based RNASeq. Illumina-based RNASeq is the most suitable experimental approach for accurate assessment of intra-host SARS-CoV-2 sequence variability–the main focus of our example application. It avoids amplification biases characteristic of PCR-based enrichment approaches such as PrimalSeq^15^–a primary methodology used to generate Illumina- and ONT-based ampliconic datasets. However, because one of the key objectives of this study is to provide freely accessible workflows for the analysis of *all types* of SARS-CoV-2 sequence data we developed procedures for handling ampliconic data as well. Thus here we will describe two distinct analytical strategies: one for Illumina-based RNASeq data and another for Amplicon data. We have also developed workflows for the analysis of ONT data including AV analysis, for consensus sequence building from variants called with any of our upstream workflows, and for processing of direct RNA sequencing data. These will be described in a separate report.

**Figure 2.**
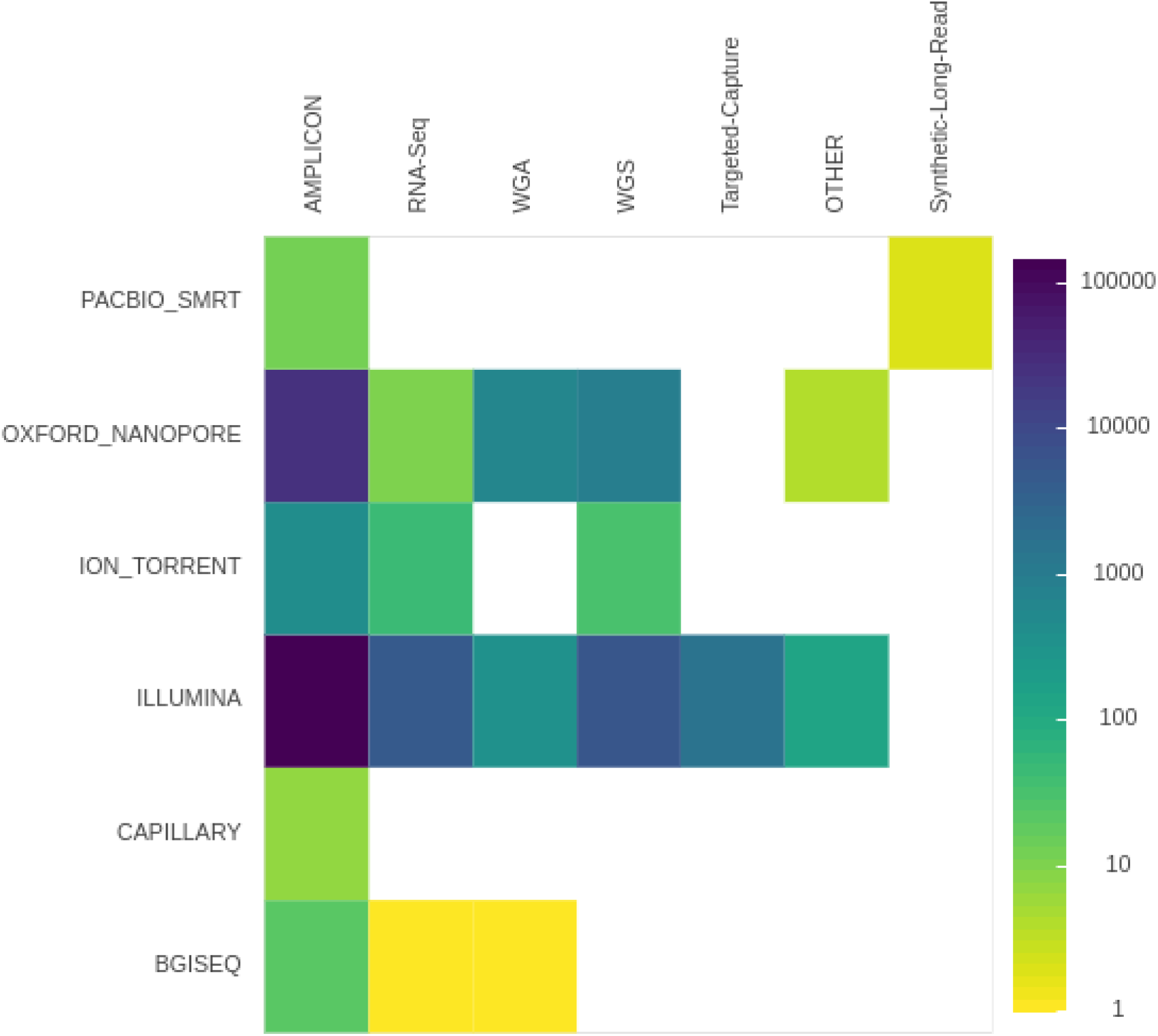
Number of SRA accession for each sequencing technology and library preparation strategy.

For each of the Illumina RNA-seq and Illumina ARTIC data generation approaches we initially identified the ten studies utilizing these approaches that yielded the largest numbers of individual read datasets (Supplementary Table 1). From these ten studies, we filtered out all those without associated publications. Because metadata for SRA datasets are generally of low quality, associated publications reveal important information about the datasets such as, for example, which ARTIC primer set versions were used or how RNA was isolated. Of datasets with associated publications we settled on two (PRJNA622837 and PRJEB37886), for subsequent analysis (see Methods). Whereas PRJNA622837 focused on the analysis of SARS-CoV-2 transmission in the Boston area of the USA^16^, PRJEB37886 is from the ongoing UK genomic surveillance effort^17^. While the entire PRJNA622837 dataset was gathered before September 2020, prior to emergence of N501Y lineages of concern (B.1.1.7, B.1.351 and P.1), the PRJEB37886 dataset contains samples isolated both before and after the emergence of these lineages. Since we were interested in determining whether signature mutations found in the N501Y lineages were detectable, and possibly detectably evolving under positive selection, in the datasets prior to the emergence of the N501Y lineages, we separated the PRJEB37886 dataset into a pre-N501Y lineage emergence dataset, hereafter called “COG-Pre” (containing accessions ERR4603708 - ERR4604210) and a post-N501Y lineage emergence dataset, hereafter called “COG-Post” (accessions ERR4859723 - ERR4861540).

We generated raw AV lists for all these datasets (Table 2) by applying the Illumina RNAseq workflow (#2 in Table 1) to the Boston dataset and the ARTIC workflow (#3 in Table 1) to the COG-Pre and COG-Post datasets. Only AVs that both occurred at an allele frequency (AF) 5% or greater, and were supported by 10 or more reads were included (see Methods). After identifying AVs we used the reporting workflow to generate a final AV summary–a single dataset listing all AVs in all samples. For each AV, the report was richly annotated, including information on per strand counts and allele frequencies for all samples, and potential functional impacts of each AV (Table 3).

**Table 3.** Allelic-variant (AV) counts pre/post filtering. AVs= number of all detected AVs; Sites = number of distinct variable sites across SARS-CoV-2 genome; Samples = number of samples in the corresponding dataset. These datasets will also be available from the Viral Beacon project at (https://covid19beacon.crg.eu/).

| Dataset | Links | AVs | Sites | Samples |
| --- | --- | --- | --- | --- |
| "Boston" | by sample<br>by variant | 9,249/8,492 | 1,027/315 | 639 |
| "COG-Pre" | by sample<br>by variant | 7,338/4,761 | 2,747/287 | 503 |
| "COG-Post" | by sample<br>by variant | 38,919/38,813 | 5,760/1,795 | 1,818 |

Exploratory analysis of AV data was then carried out in Jupyter and ObservableHQ notebooks (Table 2). For demonstration purposes we performed similar types of analysis in both Jupyter and ObservableHQ to highlight their utility. The notebooks include a number of data manipulation steps that calculate general descriptive statistics, plot the distributions of AVs across the genome and perform a variety of additional analyses that are important for contextualizing SARS-CoV-2 AVs. These additional analyses are briefly summarized below.

Any set of AV calls unavoidably includes erroneous calls that need to be filtered out. We assumed that a fraction of AVs with low frequencies are random errors, modeled by a simple Poisson distribution with per-site error rate λ. We then tabulated, for each position in the genome, the number of samples that contained an AV with 0.05 ≤ AF ≤ 0.5, inferred λ using a closed form ML estimator (the mean of per-base counts), and plotted the observed number of genome positions with *N* = 0,1,2… AV (Fig. S1). In all three datasets the point where the predicted Poisson distribution clearly diverged from the observed distribution (*N* = 3 for the Boston dataset, and *N* = 2 for the COG-Pre and COG-Post datasets) could be taken as the error-vs-real threshold (Fig. S1). Applying these thresholds to the data reduced AV counts as shown in Table 3 after the “/” symbol.

When considering AVs with all AFs, the dominant patterns of co-occurrence were clade-segregating sites in the data, e.g. high frequency AVs that exist in strong linkage disequilibrium (e.g. the 241/3037/14408/24403/25563 set seen as thick vertical lines in Fig. 3A). A more interesting pattern was observed when we restricted our attention only to relatively common low frequency AVs (Fig. 3B) among which there were several groups that co-occured in multiple samples (all exclusively at low frequencies). A cluster of eight low frequency AVs was identified in eight samples (Table 4; the probability of this occurring by chance is < 10^−8^). No similar low AF clusters were detected in either the COG-pre or COG-post datasets, but a cluster of two medium AF AVs (9096:C→T, 29692:G→T) co-occurred three times (expected *p* < 0.01; not shown).

**Figure 3.**
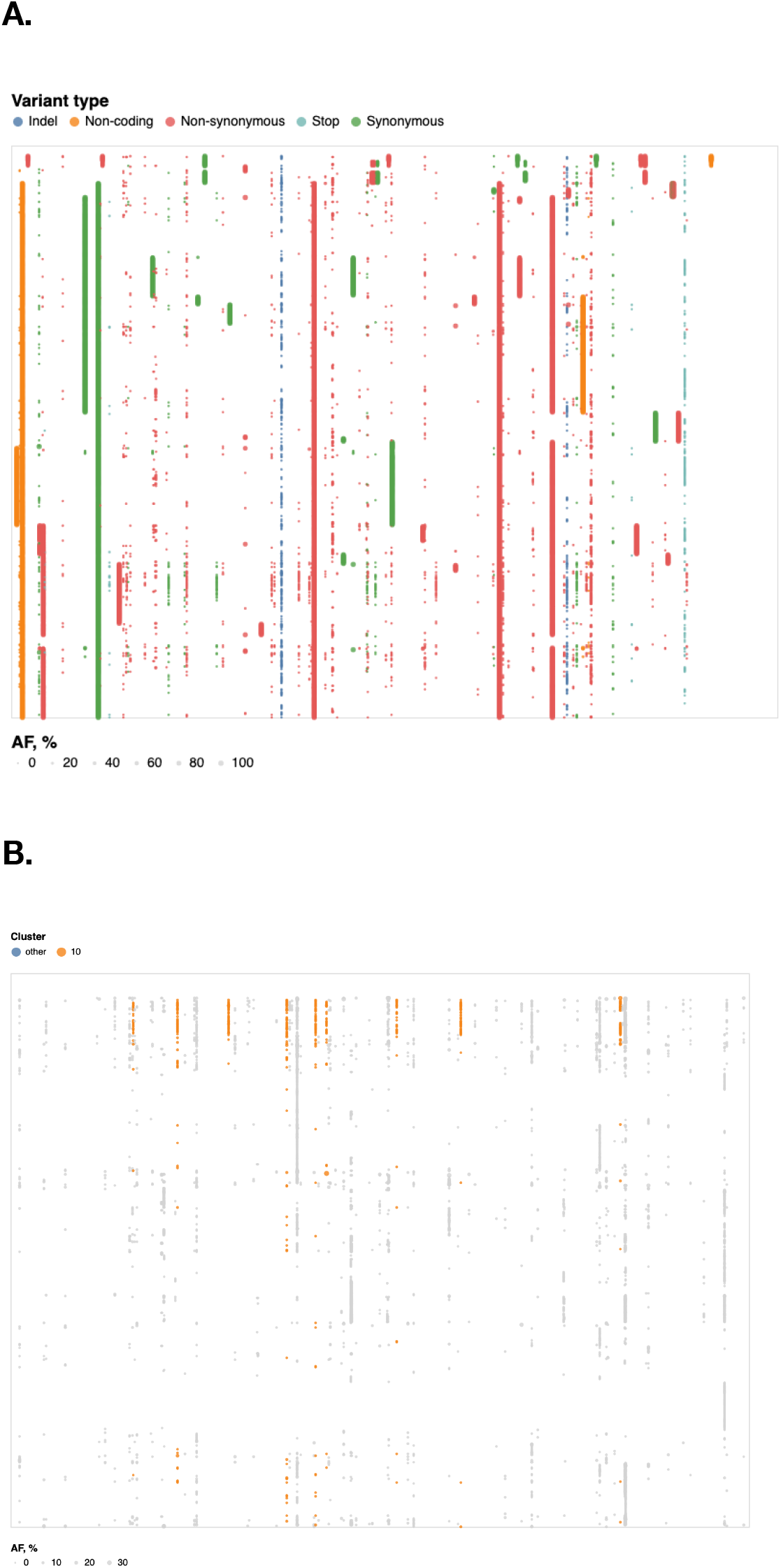
Dot plot of all allelic-variants (AV) across samples in Boston dataset. X-axis: genome position, Y-axis: Samples, colors correspond to functional classes of AVs. Samples are arranged by hierarchical clustering using cosine distances on mean allele frequencies of all AVs. **A**. Dot-plot of all allelic variants in the “Boston” dataset; rows – samples, columns – genomic coordinates; samples are arranged by hierarchical clustering. Limited to variants that occur in at least 4 samples. **B**. Dot-plot of observed variants in the “Boston” dataset; restricted to variants that appear only at AF≤10% and occur in at least 4 samples each. Variants are partitioned into 10 clusters, using K-medoids using the Hamming distance on AF vectors; the cluster with 8 variants is highlighted in orange.Interactive version is at https://observablehq.com/@spond/intrahost-variant-exploration-landing

**Table 4.** Eight low frequency allelic-variants co-occurred in 8 samples.

| Nucleotide Variant | Sample count | Effect |
| --- | --- | --- |
| 4,338:C→T | 29 | nsp3/S540F |
| 6,604:A→G | 54 | nsp3/L1295 |
| 9,535:C→T | 37 | nsp4/T327 |
| 12,413:A→C | 63 | nsp8/N108H |
| 13,755:A→C | 53 | RdRp/R105S |
| 14,304:A→C | 30 | RdRp/K288N |
| 17,934:C→A | 30 | helicase/T566 |
| 20,716:A→T | 37 | MethTr/M20L |
| 26,433:A→C | 35 | E/K63N |

Any newly identified AV set should be compared against a list of sites with known biological significance. In our analysis we selected two such site lists: (1) the signature or defining mutation sites found in the variants of concern (B.1.1.7, B.1.351, P.1 and others referred to hereafter as VOC sites); and (2) genomic sites detectably evolving under positive or negative selection in the global SARS-CoV-2 dataset (referred to hereafter as selected sites).

The emergence of VOC starting with the B.1.1.7 lineage in the UK, raised intriguing questions about the genesis of this lineage, and a hypothesis that the variant arose in a chronically infected immunocompromised host^18^. We were interested in how many of the clade defining mutations^19^ were detectable at sub-consensus allele frequencies. Specifically, we analyzed the overlap between our data and five distinct mutation sets: B.1.1.7, P.1, B.1.351, A.23.1 as well as receptor-binding domain mutations identified by Greaney *et al.^20^* (Fig. 4). Very few N501Y lineage mutations are detectable in pre B. 1.1.7 datasets (Fig. 4). The L18F mutation (P.1, gene *S*) is present in ~0.5% of Boston samples. It is completely absent in “COG-Pre”, while in “COG-Post” it reaches fixation in ~40% of samples. On the other hand the population frequency of E92K (P.1, gene *S*) stays constant for “COG-Pre” and “COG-Post” samples at ~3%. Only two of the mutations reported by Greaney *et al*.^20^ are observed at appreciable frequency.

**Figure 4.**
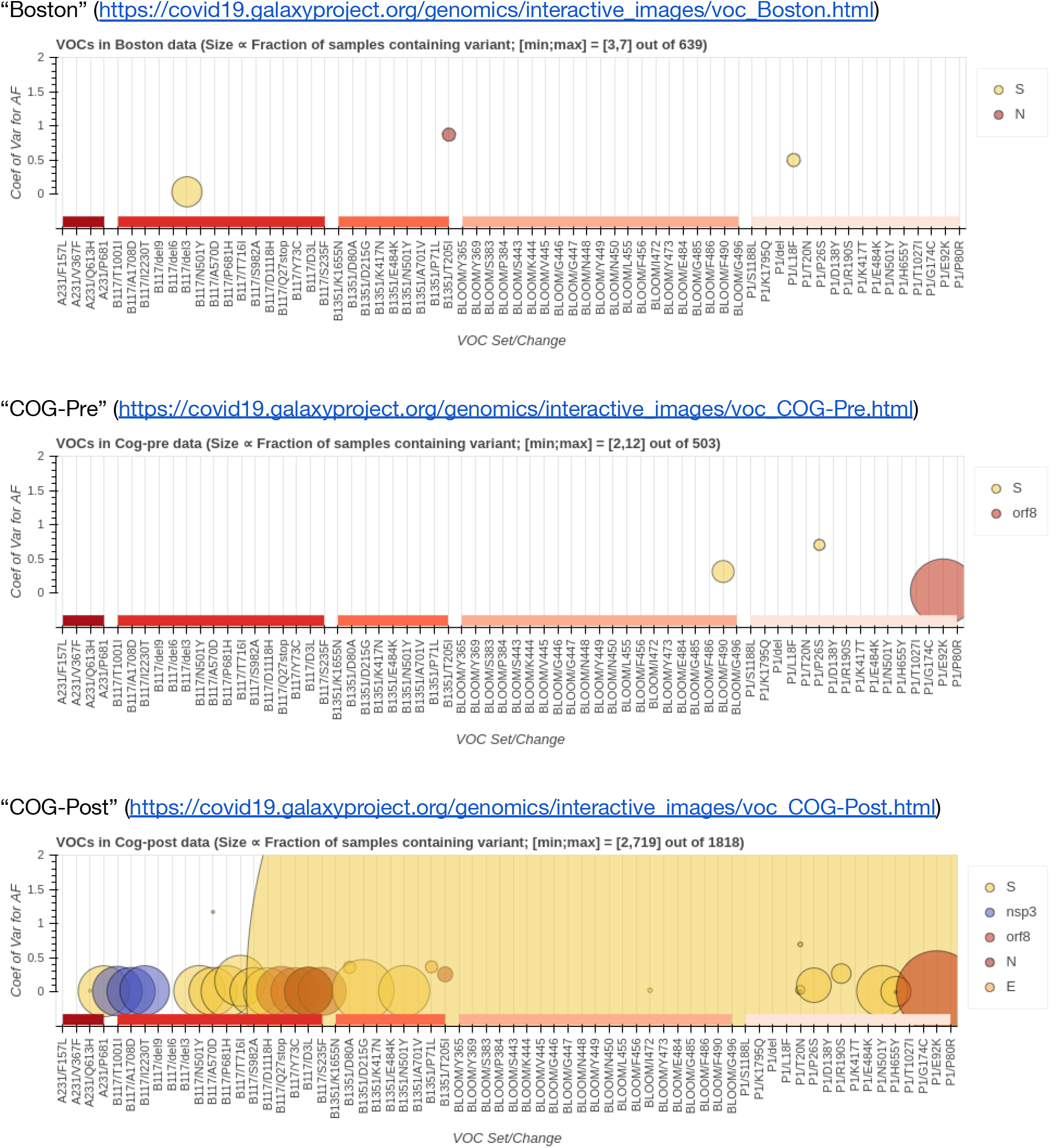
Intersection between allelic-variants (AV) reported here with AVs of concern (VOC). Big blob in “COG-Post” dataset corresponds to L18F change in gene S. Size of markers ∞ fraction of samples containing variant. [min;max] - maximum and minimum counts of samples containing variants shown in this figure. E.g., in “Boston” the largest marker corresponds to an AV shared by 7 samples, and the smallest by 3 samples.

The selected sites list is continuously updated by the DataMonkey team using GISAID data as it accumulates (http://covid19.datamonkey.org). This list included all SARS-CoV-2 codon sites identified with FEL^21^ and MEME^22^ methods to be evolving under positive or negative selection with a *p*≤0.0001 significance cutoff in all GISAID datasets as of Feb 01, 2021. Because selection analyses identify codons (not individual genome positions) responsible for potential selective amino acid changes, we considered all non-synonymous nucleotide substitutions with AFs< 80% that fell within the boundaries of codons with the signature of selection. There were two, three, and ten AVs overlapping with codons under selection in the Boston, COG-Pre, and COG-Pos” datasets, respectively (Table 5). The two sites in the Boston dataset are below the consensus (<50%) frequency. The identification of low frequency AVs at codon sites displaying evidence of positive selection could provide an early indication of AVs that are adaptive mutations and therefore warrant closer monitoring as the pandemic unfolds.

**Table 5.** Allelic-variants (AVs) with maximum allele frequency < 80% overlapping with codons under selection. % samp = fraction of samples containing a given AV in each dataset

| POS | R | A | Gene | AA | % samp | mean | min | max | codon |
| --- | --- | --- | --- | --- | --- | --- | --- | --- | --- |
| "Boston" |  |  |  |  |  |  |  |  |  |
| 25,842 | A | C | orf3a | T151P | 6.259 | 0.083 | 0.051 | 0.180 | ACT |
| 22,254 | A | G | S | I231M | 2.034 | 0.066 | 0.050 | 0.102 | ATA |
| "COG-Pre" |  |  |  |  |  |  |  |  |  |
| 21,637 | C | T | S | P26S | 0.397 | 0.520 | 0.260 | 0.779 | CCT |
| 22,343 | G | T | S | G261V | 0.397 | 0.425 | 0.091 | 0.758 | GGT |
| 28,825 | C | T | N | R185C | 0.397 | 0.619 | 0.528 | 0.709 | CGT |
| "COG-Post" |  |  |  |  |  |  |  |  |  |
| 1,463 | G | A | nsp2 | G220D | 0.715 | 0.737 | 0.669 | 0.767 | GGT |
| 22,343 | G | T | S | G261V | 0.660 | 0.703 | 0.672 | 0.737 | GGT |
| 25,217 | G | T | S | G1219<br>V | 0.330 | 0.752 | 0.746 | 0.756 | GGT |
| 21,845 | C | T | S | T95I | 0.275 | 0.466 | 0.069 | 0.731 | ACT |
| 29,252 | C | T | N | S327L | 0.220 | 0.196 | 0.083 | 0.359 | TCG |
| 29,170 | C | T | N | H300Y | 0.165 | 0.314 | 0.076 | 0.735 | CAT |
| 4,441 | G | T | nsp3 | V575L | 0.110 | 0.725 | 0.712 | 0.739 | GTG |
| 21,829 | G | T | S | V90F | 0.110 | 0.508 | 0.242 | 0.774 | GTT |
| 28,394 | G | A | N | R41Q | 0.110 | 0.318 | 0.054 | 0.582 | CGG |
| 29,446 | G | T | N | V392L | 0.110 | 0.770 | 0.765 | 0.774 | GTG |

### Continuous analysis of pandemic data with the Galaxy API

Genome surveillance projects at national levels like COG-UK produce sequencing data for unprecedented numbers of samples. To demonstrate that our system can satisfy the analysis needs of such projects, we are performing near real-time analysis of COG-UK sequencing data as it is being submitted to the European Sequence Archive (ENA). We set up an automated analysis system that runs our ARTIC AV-calling workflow and the reporting workflow programmatically on all new COG-UK ARTIC paired-end data via Galaxy’s openly accessible API. The system also handles data organization into Galaxy histories and exports of resulting datasets. The datasets analyzed at the time of submission of this paper are listed in Table S2. Table S2 also provides a link to a location containing a continuously updated version of this table. These datasets will also be available from the Viral Beacon project at (https://covid19beacon.crg.eu/).

### The bottom line

The system we describe here can be used right now by anyone in the world with an internet connection simply by pointing a standard web-browser to usegalaxy.{org|eu|org.au}. No software installation or hardware configuration is necessary. Our platform is powered by global public research computing infrastructure, which is able to sustain large analyses at scale. While we believe that our workflows and notebooks are good starting points for comprehensive analyses of AVs, they are not set in stone. In fact, we expect that the community will improve them via continuous updates and modifications. To facilitate this process the workflows are readily available from the global repositories, https://dockstore.org and https://workflowhub.eu. They can be modified by anyone using the Galaxy workflow editor and adapted to individual research needs by replacing or adding tools. Similarly, Jupyter and ObservableHQ notebooks can be used as-is or further enhanced.

The emergence of SARS-CoV-2 variants of concern in the UK, Brazil and South Africa is probably more indicative of the monitoring infrastructures of these countries than it is of some unique characteristic(s) of their respective COVID-19 epidemics that specifically favoured the emergence of these variants. Emergence of divergent lineages is an expected general feature of endemic RNA pathogens, and it is likely that many other potential variants of concern remain undetected. While our system is specifically designed to encourage collaborative worldwide genomic surveillance to rapidly identify and respond to such variants, its use of raw read data rather than assembled genomes goes a step beyond current surveillance efforts. Specifically it enables the coordinated worldwide surveillance of intra-patient minor AV frequencies: a distinction that could yield decisive early-warnings of epidemiological conditions conducive to the emergence of variants with altered pathogenicity, vaccine sensitivity or resistance. One lesson of the COVID-19 pandemic has been that uncoordinated efforts to gather such information are not much better than gathering no information at all. Another much harsher lesson has been that failing to effectively use such information could cost hundreds of thousands of lives.

## Methods

### Analysis workflows

Analysis workflows are listed in Table 1. The two Illumina RNASeq workflows (#1 and #2 in Table 1) perform read mapping with `bwa-mem` and `bowtie2`, respectively, followed by sensitive allelic-variant (AV) calling across a wide range of AFs with `lofreq` (see AV calling section below).

The workflow for Illumina-based ARTIC data (#3 in Table 1) builds on the RNASeq workflow for paired-end data using the same steps for mapping and AV calling, but adds extra logic operators for trimming ARTIC primer sequences off reads with the `ivar` package. In addition, this workflow uses `ivar` also to identify amplicons affected by ARTIC primer-binding site mutations and excludes reads derived from such “tainted” amplicons when calculating AFs of other AVs.

The workflow for ONT-sequenced ARTIC data is modeled after the alignment/AV-calling steps of the ARTIC pipeline (https://artic.readthedocs.io/). It performs, essentially, the same steps as that pipeline’s `minion` command, i.e. read mapping with `minimap2` and AV calling with `medaka`. Like the Illumina ARTIC workflow it uses `ivar` for primer trimming. Since ONT-sequenced reads have a much higher error rate than Illumina-sequenced reads and are therefore plagued more by false-positive AV calls, this workflow makes no attempt to handle amplicons affected by potential primer-binding site mutations.

All four workflows use `SnpEff`, specifically its `4.5covid19` version, for AV annotation.

The fifth workflow (Reporting) takes a table of AVs produced by any of the other four workflows and generates a list of AVs by Samples and by Variant. For an example see [here](https://github.com/galaxyproject/SARS-CoV-2/raw/master/data/var/).

Workflows default to requiring an AF ≥ 0.05 and AV-supporting reads of ≥ 10 (these and all other parameters can be easily changed by the user). For an AV to be listed in the reports it must surpass these thresholds in at least one sample of the respective dataset. We estimate that for AV calls with an AF ≥ 0.05 our analyses have a false-positive rate of < 15% for both Illumina RNAseq and Illumina ARTIC data, while the true-positive rate of calling such low-frequency AVs is ~80% and approaches 100% for AVs with an AF ≥0.15. This estimate is based on an initial application of the Illumina RNAseq and Illumina ARTIC workflows to two samples for which data of both types had been obtained at the virology department of the University of Freiburg and the assumption that AVs supported by both sets of sequencing data are true AVs. The second threshold of 10 AV-supporting reads is applied to ensure that calculated AFs are sufficiently precise for all AVs.

### Selection of the AV caller

The development of modern genomic tools and formats have been driven by large collaborative initiatives such as 1,000 Genomes, GTEx and others. As a result, the majority of current AV callers have been originally designed for diploid genomes of human or model organisms where discrete AFs are expected. Bacterial and viral samples are fundamentally different. They are represented by mixtures of multiple haploid genomes where the frequencies of individual AVs are treated as a continuous variable. This renders many existing AV calling tools unsuitable for microbial and viral studies unless one is looking for fixed AVs. However, recent advances in cancer genomics have prompted developments of somatic AV calling approaches that do not require normal ploidy assumptions and can be used in the analysis of either samples with chromosomal malformations or circulating tumor cells. The latter situation is essentially identical to viral or bacterial resequencing scenarios. As a result of these developments the current set of AV callers appropriate for microbial studies includes updated versions of “legacy” tools (`FreeBayes`^23^ and `mutect2`^24^) as well as dedicated packages (`Breseq`^25^, `SNVer`^26^, and `lofreq`^27^). To assess the applicability of these tools we first considered factors related to their long-term sustainability, such as the health of the codebase as indicated by the number of code updates, contributors and releases as well as the number of citations. After initial testing we settled on three AV callers: `FreeBayes`, `mutect2`, and `lofreq` (`Breseq`’s new “polymorphism mode” was in an experimental state at the time of testing. SNVer is no longer actively maintained). `FreeBayes` contains a mode specifically designed for finding sites with continuous AFs; `Mutect2` features a so-called mitochondrial mode, and `lofreq` was specifically designed for microbial sequence analysis.

We are seeking to be able to detect AVs with frequencies around the NGS detection threshold of ~ 1-5%^27^. In order to achieve this goal we selected a test dataset, which is distinct from data used in recent method comparisons^28,29^. These data originate from a duplex sequencing experiment recently performed by our group 25. In this dataset a population of *E. coli* cells transformed with pBR322 is maintained in a turbidostat culture for an extensive period of time. Adaptive changes accumulated within the plasmid are then revealed with duplex sequencing^30^. Duplex sequencing allows identification of AVs at very low frequencies. This is achieved by first tagging both ends of DNA fragments to be sequenced with unique barcodes and subjecting them to paired-end sequencing. After sequencing, read pairs containing identical barcodes are assembled into families. This procedure allows one to reliably separate errors introduced during library preparation and/or sequencing (present in some but not all members of a read family) from true AVs (present in all members of a read family derived from both strands).

For the following analysis we selected two data points from Ref.^30^: one corresponding to the beginning of the experiment (s0) and the other to the end (s5). The first sample is expected to be nearly clonal with no variation, while the latter contains a number of adaptive changes with frequencies around 1%. We aligned duplex consensus sequences (DCS) against pBR322. We then walked through read alignments to produce counts of non-reference bases at each position (Fig. S6).

Because all differences identified this way are derived from DCS reads they are a reasonable approximation for a “true” set of AVs. s0 and s5 contained 38 and 78 variable sites with at least two alternative counts, respectively (among 4,361 bases on pBR322) of which 27 were shared. We then turned our attention to the set of sites that were determined by Mei et al. to be under positive selection (sites 3,029, 3,030, 3,031, 3,032, 3,033, 3,034, 3,035, and 3,118). Changes at these sites increase the number of plasmid genomes per cell. Sample s0 does not contain alternative bases at any of these sites. Results of the application of the three AV callers with different parameter settings (shown in Table S1) are summarized in Fig. S7.

Overall `lofreq` performed the best followed by `mutect2` and `FreeBayes` (contrast “Truth” with “nf” and “def” in Fig. S7). The main disadvantage of `mutect2` is in its handling of multiallelic sites (e.g., 3,033 and 3,118) where multiple alternative bases exist. At these sites `mutect2` outputs alternative counts for only one of the AVs; the one with highest counts. This is why at site 3,118 A and T counts are identical. Given these results we decided to use `lofreq` for the main analysis of the data.

## Acknowledgements

The authors are grateful to the broader Galaxy community for their support and software development efforts. This work is funded by NIH Grants U41 HG006620 and NSF ABI Grant 1661497. Usegalaxy.eu is supported by the German Federal Ministry of Education and Research grants 031L0101C and de.NBI-epi to BG. Galaxy and HyPhy integration is supported by NIH grant R01 AI134384 to AN. Usegalaxy.org.au is supported by Bioplatforms Australia and the Australian Research Data Commons through funding from the Australian Government National Collaborative Research Infrastructure Strategy. hyphy.org development team is supported by NIH grant R01GM093939. Usegalaxy.be is supported by the Research Foundation-Flanders (FWO) grant I002919N and the Flemish Supercomputer Center (VSC). The funders had no role in study design, data collection and analysis, decision to publish, or preparation of the manuscript.

---

**Table S1.** Parameters used for comparing performance of allelic-variant callers

| Caller | Command line | Figure 2 label |
| --- | --- | --- |
| mutect2 | --mitochondria-mode true | m |
| mutect2 | default | m_noM |
| mutect2 | --mitochondria-mode true<br>--f1r2-max-depth 1000000 | m_md_inf |
| mutect2 | --mitochondria-mode true<br>--f1r2-max-depth 1000000<br>-max-af 1 | m_md_inf_max_af1 |
| freebayes | --haplotype-length 0<br>--min-alternate-fraction 0.001<br>--min-alternate-count 1<br>--pooled-continuous --ploidy 1 | hl-0_maf-001_pc |
| freebayes | -min-alternate-fraction 0.001<br>--pooled-continuous --ploidy 1 | maf-001_pc |
| lofreq | --no-default-filter | nf |
| lofreq | default | def |

**Table S2.** Links to Galaxy histories containing COG-UK reanalysis results. A continuously updated list can be accessed https://github.com/galaxyproject/SARS-CoV-2/tree/master/data/cog-uk-tracking.These datasets will also be available from the Viral Beacon project at (https://covid19beacon.crg.eu/).

| Galaxy histories | contained ENA IDs | AV history (with VCF of all called variants) | variant report history (with tabular reports of all variants) |
| --- | --- | --- | --- |
| COG-UK NT1659949O | ERR5379175-ERR5379540 | https://usegalaxy.eu/histories/view?id=a70e27d468a29519 | - |
| COG-UK NT1659948N | ERR5378946-ERR5379174 | https://usegalaxy.eu/histories/view?id=d67968ed5fe74586 | https://usegalaxy.eu/histories/view?id=762cd32fdd59b3ef |
| COG-UK NT1659945K | ERR5352103-ERR5352455 | https://usegalaxy.eu/histories/view?id=97050f0c7217eb53 | https://usegalaxy.eu/histories/view?id=1bf724bcb1a6bce2 |
| COG-UK NT1659944J | ERR5351745-ERR5352102 | https://usegalaxy.eu/histories/view?id=c5156eda4b3c5e24 | https://usegalaxy.eu/histories/view?id=55823ef9c77a410d |
| COG-UK NT1659891N | ERR5378314-ERR5378575 | https://usegalaxy.eu/histories/view?id=c25d0f298ac2ef34 | https://usegalaxy.eu/histories/view?id=d30fa7c11c245bd7 |
| COG-UK NT1659885P | ERR5352718-ERR5375533 | https://usegalaxy.eu/histories/view?id=8f829af16c834443 | https://usegalaxy.eu/histories/view?id=df02b799e2c42a7c |
| COG-UK NT1660206U | ERR5382276-ERR5382582 | https://usegalaxy.eu/histories/view?id=484f288640bdbbc4 | https://usegalaxy.eu/histories/view?id=612b0916712b673a |
| COG-UK NT1660207V | ERR5382583-ERR5382926 | https://usegalaxy.eu/histories/view?id=23191ca01f1766fd | https://usegalaxy.eu/histories/view?id=1c870e4766af57fa |
| COG-UK NT1660170C | ERR5379541-ERR5379801 | https://usegalaxy.eu/histories/view?id=216df75b20548363 | https://usegalaxy.eu/histories/view?id=996170468f48ad5e |
| COG-UK NT1660168I | ERR5380719-ERR5381079 | https://usegalaxy.eu/histories/view?id=929888a678e25678 | https://usegalaxy.eu/histories/view?id=a5371f3d76143ee6 |
| COG-UK NT1660200O | ERR5380509-ERR5380718 | https://usegalaxy.eu/histories/view?id=5d19cd66540f3b4f | https://usegalaxy.eu/histories/view?id=2d5595a60fe335b8 |
| COG-UK NT1660185J | ERR5381350-ERR5381634 | https://usegalaxy.eu/histories/view?id=761b945d604a584c | https://usegalaxy.eu/histories/view?id=0bb2222cb26f62fb |
| COG-UK NT1660171D | ERR5379802-ERR5380132 | https://usegalaxy.eu/histories/view?id=ee5d68ddc0c16766 | https://usegalaxy.eu/histories/view?id=b7435d647332c545 |
| COG-UK NT1660086H | ERR5378576-ERR5378945 | https://usegalaxy.eu/histories/view?id=bdfae26e713d1461 | https://usegalaxy.eu/histories/view?id=7586ad8f459dfbdc |
| COG-UK NT1660169J | ERR5381080-ERR5381349 | https://usegalaxy.eu/histories/view?id=02dd0e471db55821 | https://usegalaxy.eu/histories/view?id=22410c20518125b3 |
| COG-UK NT1660201P | ERR5380133-ERR5380314 | https://usegalaxy.eu/histories/view?id=f584aa738ccbde76 | - |
| COG-UK NT1660199P | ERR5381954-ERR5382275 | https://usegalaxy.eu/histories/view?id=ee1e4a8c859879dd | - |
| COG-UK NT1660186K | ERR5381635-ERR5381953 | https://usegalaxy.eu/histories/view?id=9dab59078524857b | - |

**Figure S1.**
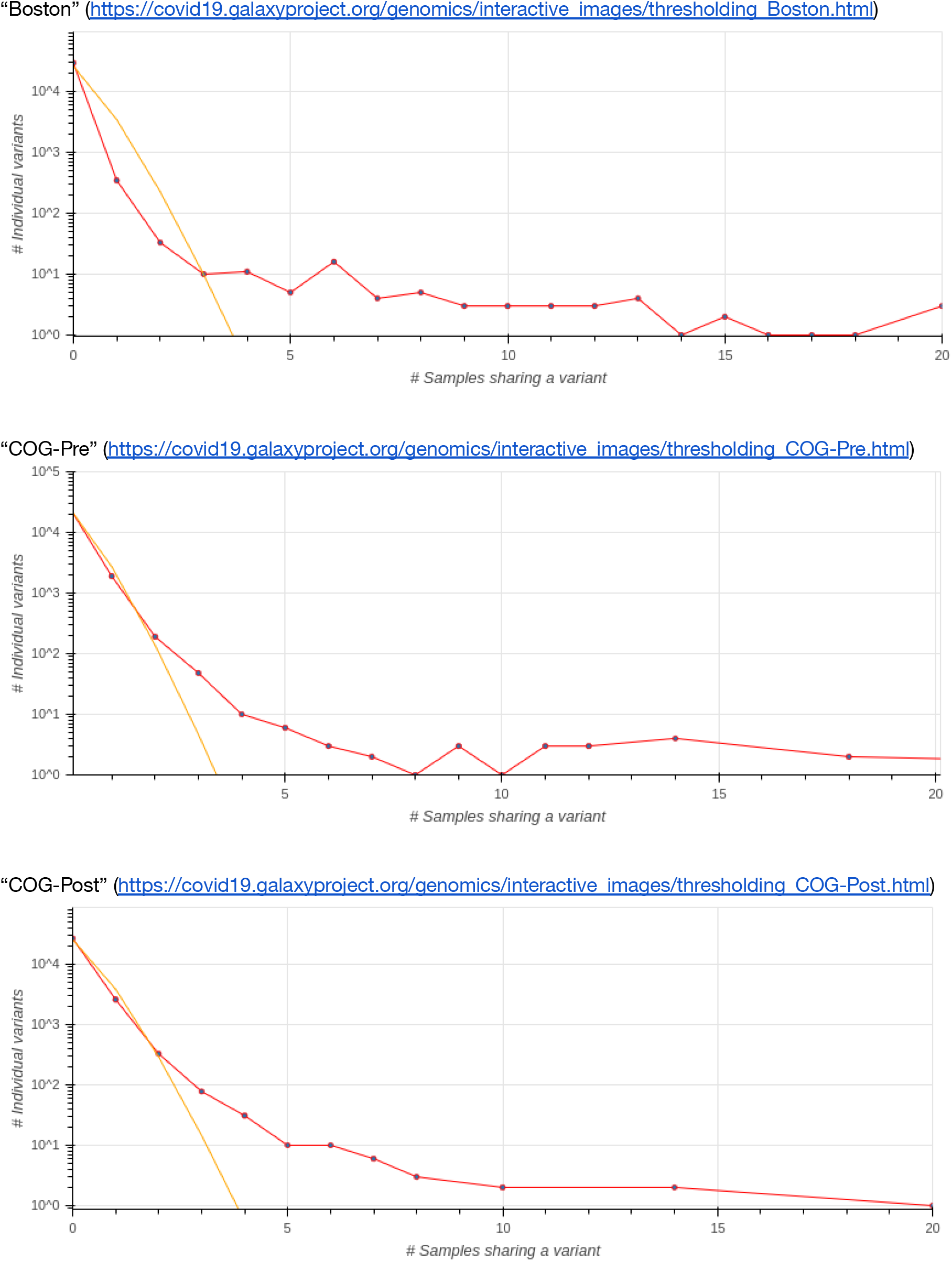
Observed (red) versus predicted (orange) counts of samples sharing *N=0, 1, 2*,… varinants as a function of allelic-variant number for each dataset.The intersection of the lines gives the cutoff that was applied to each dataset.

**Figure S2.**
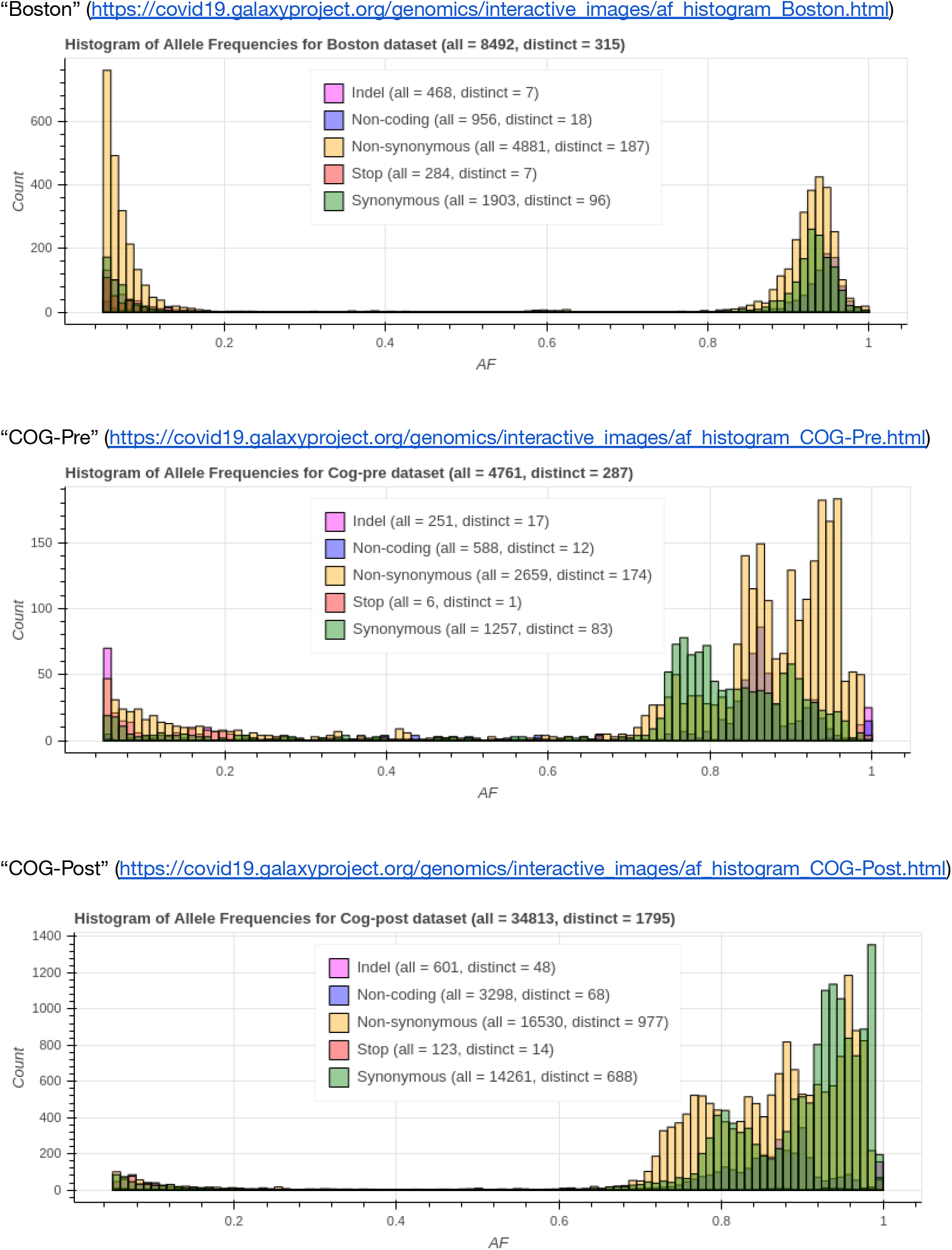
Distribution of allele frequencies for different types of substitutions.

**Figure S3.**
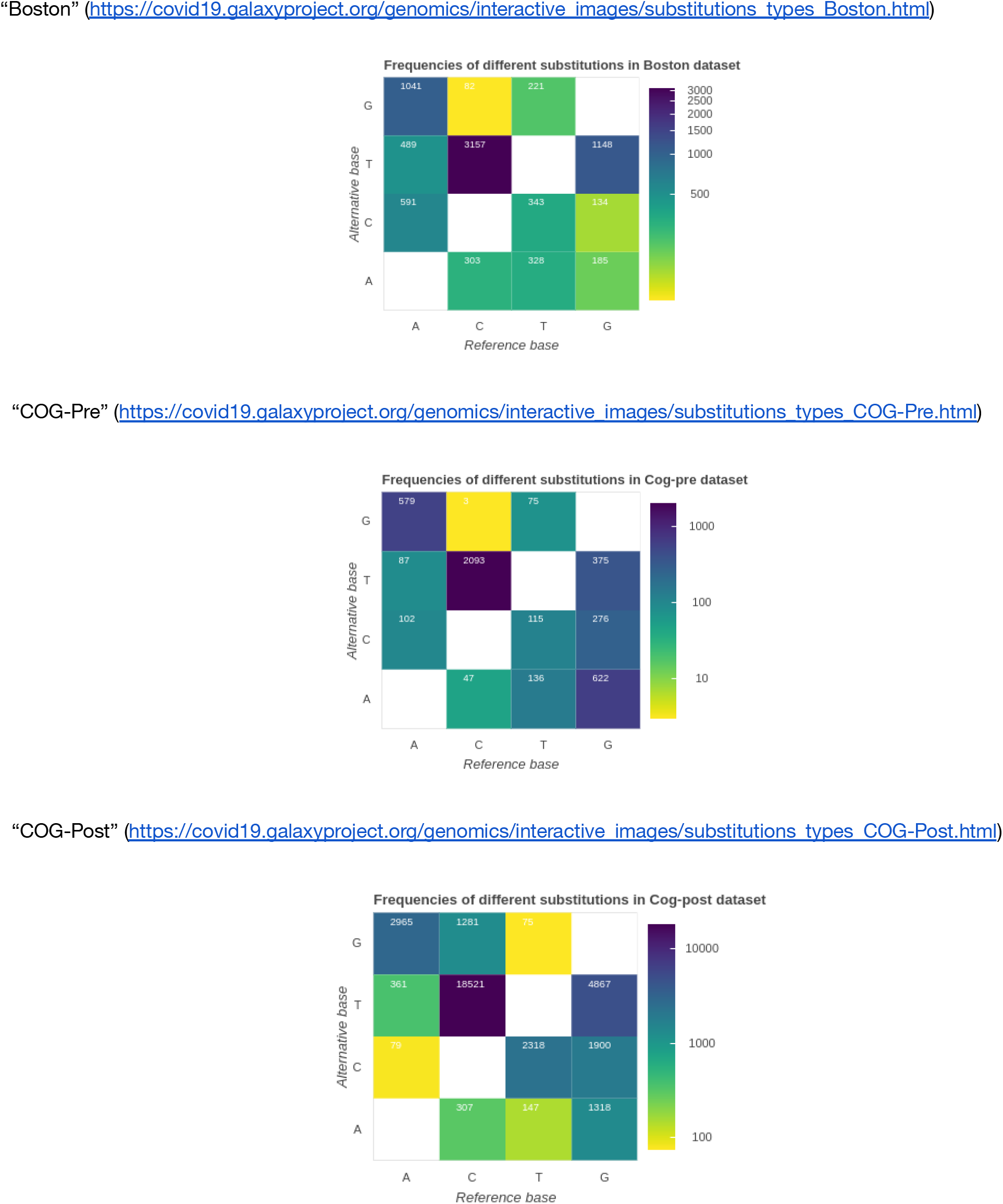
Types and counts of single nucleotide substitutions.

**Figure S4.**
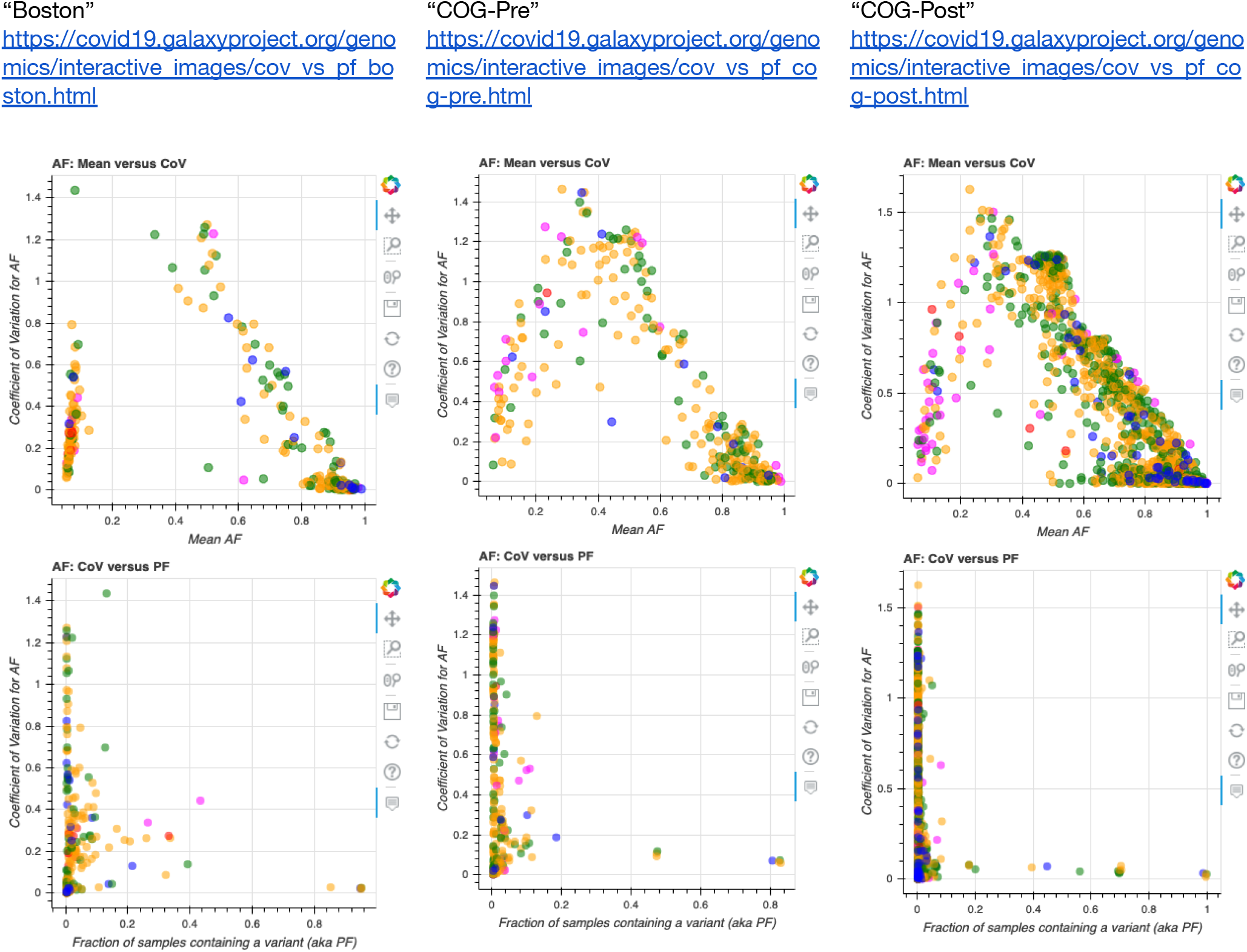
Relationship between population frequency and alternative allele frequency in the three datasets. CoV = coefficient of variation. PF = population frequency (e.g., how many samples in the dataset share a given variant). Points with high CoV have large spread of allele frequencies.

**Figure S5.**
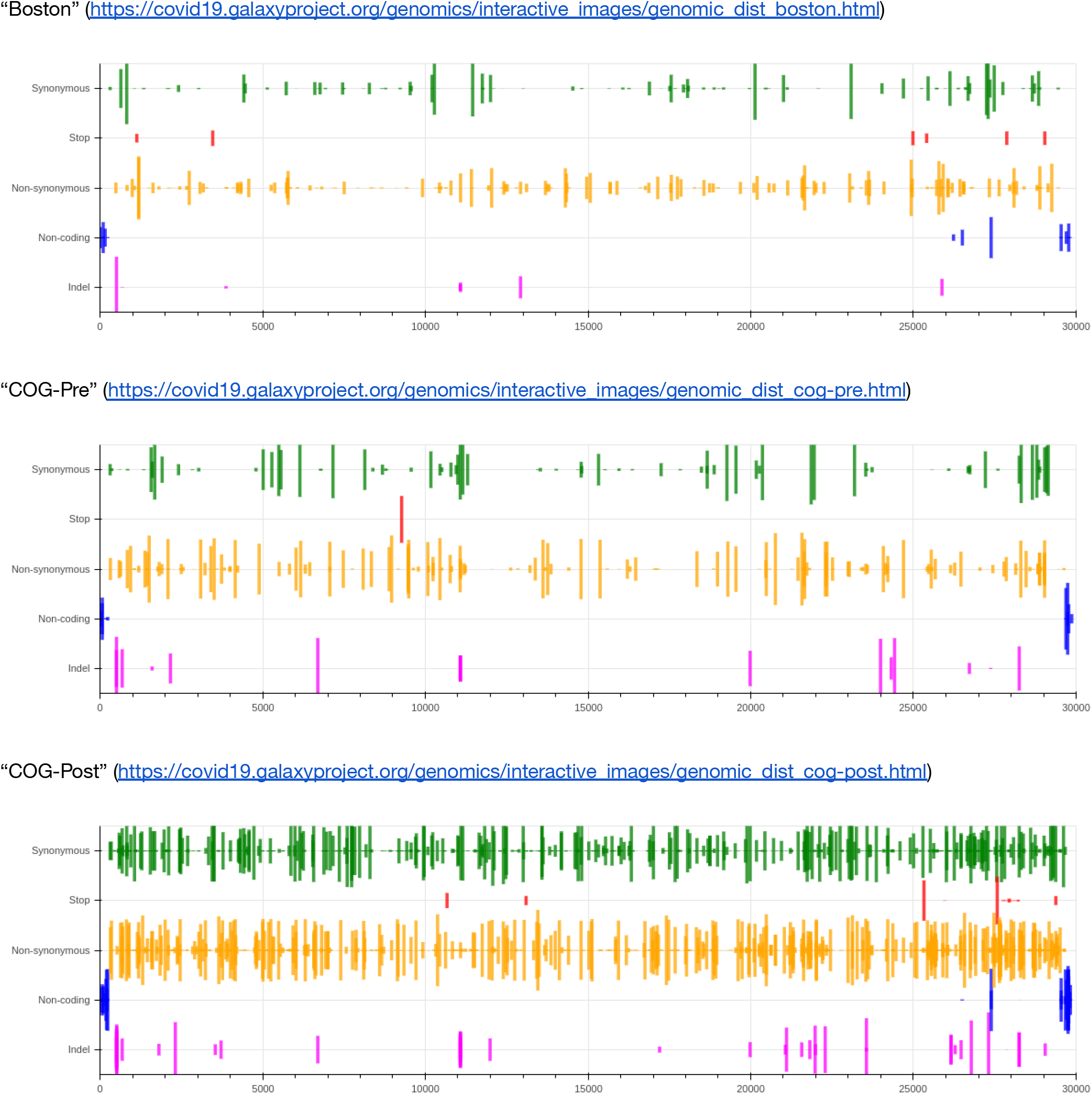
Distribution of allelic variants across the genome. Height of each bar is proportional to the coefficient of variation for alternative allele frequency.

**Figure S6.**
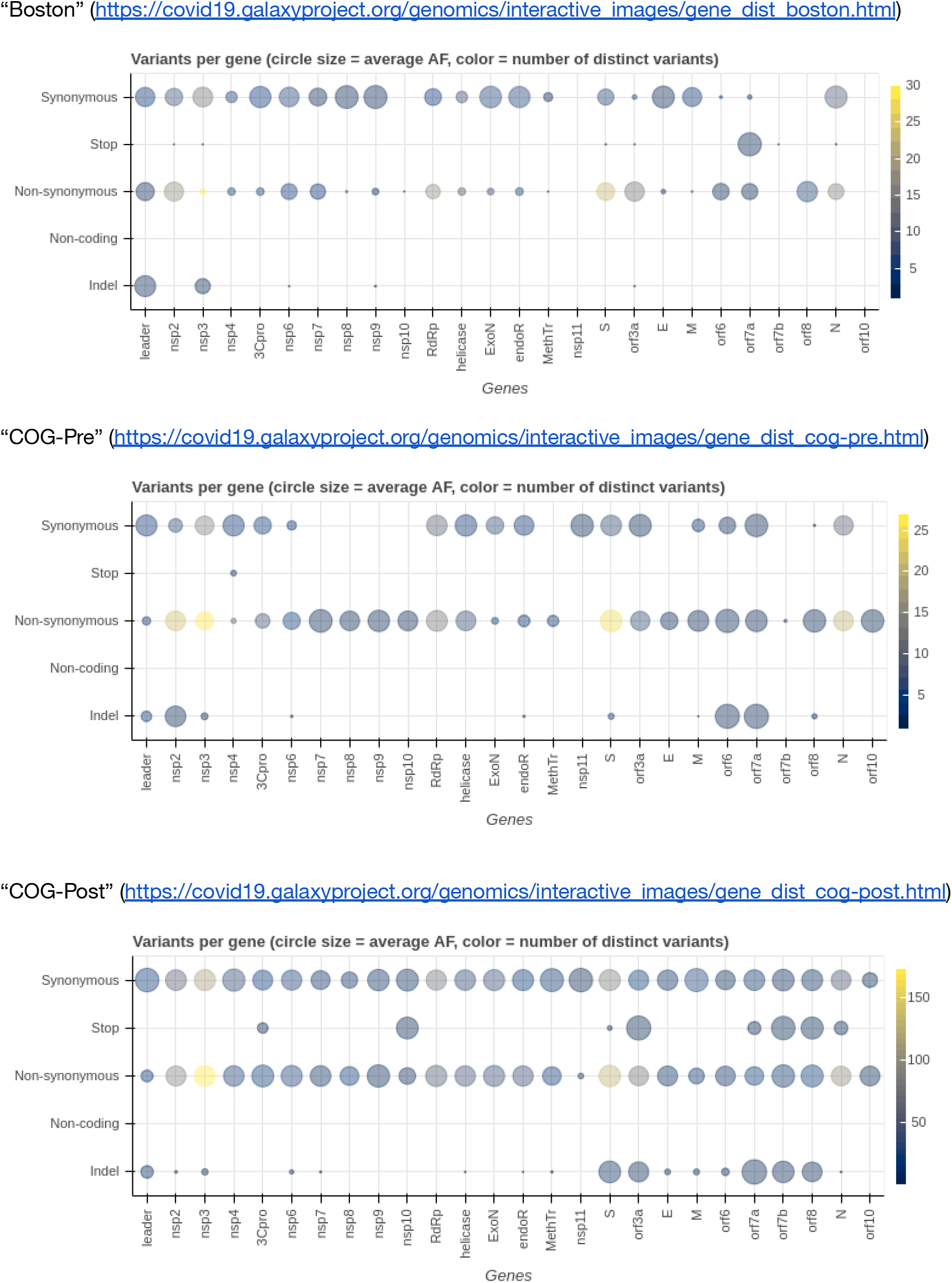
Allelic variant counts per gene.

**Figure S7.**
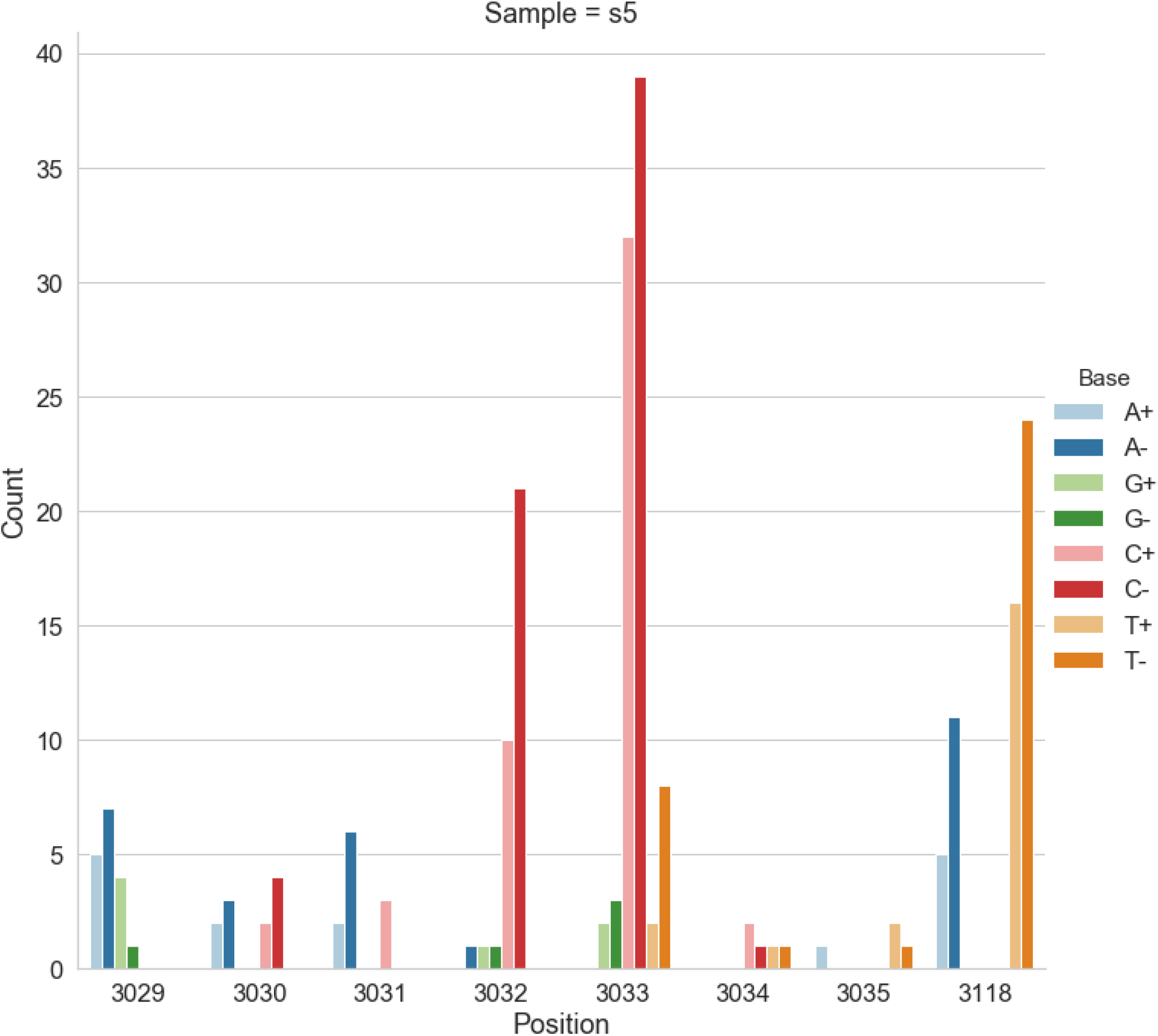
Counts of alternative bases at eight variable locations within pBR322.

**Figure S8.**
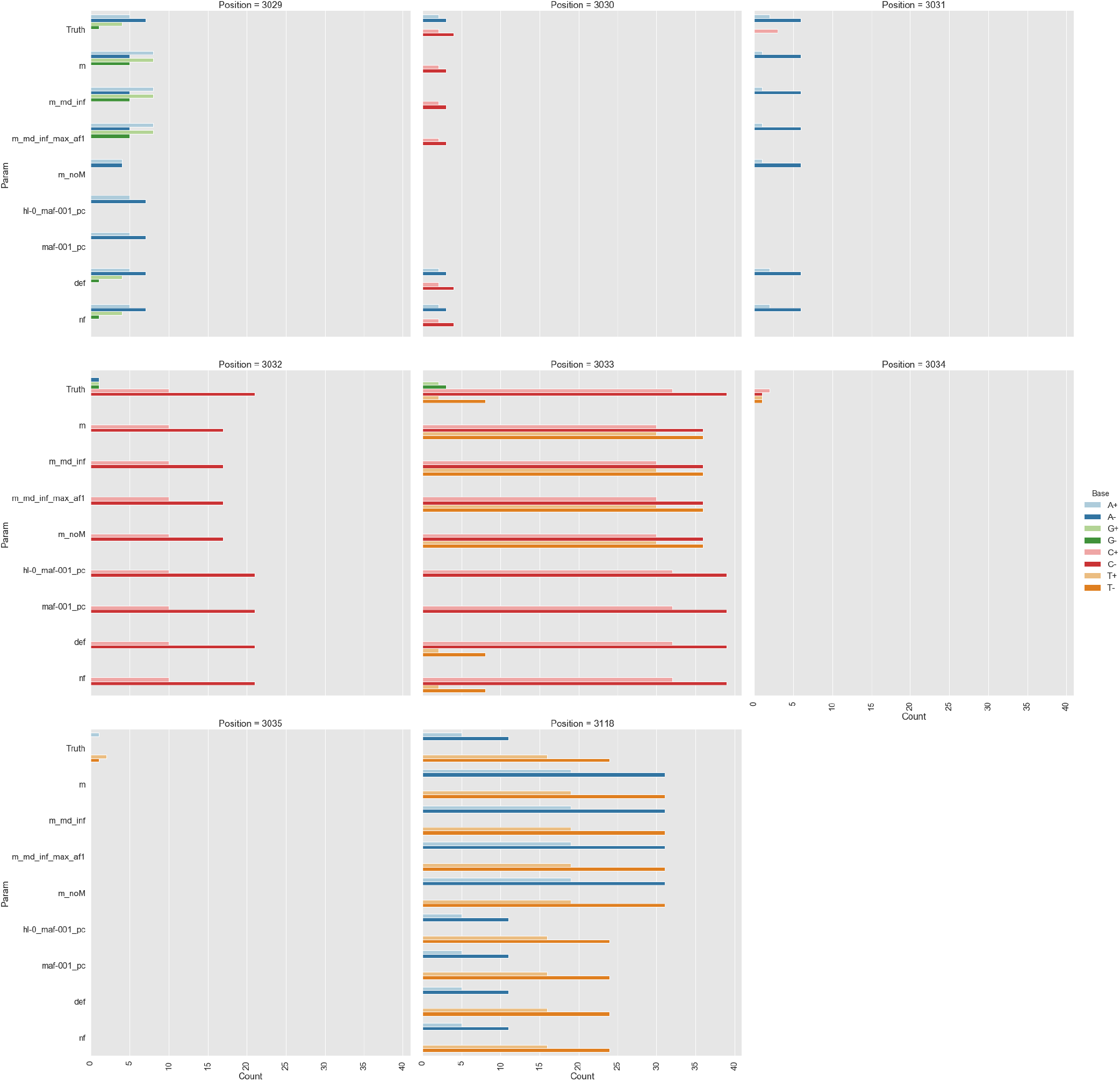
Calls made by `mutect2`, `freebayes`, and `lofreq`. For explanation of x-axis labels see Table S1.

## References

1. Baker, D. et al. No more business as usual: Agile and effective responses to emerging pathogen threats require open data and open analytics. PLoS Pathog. 16, e1008643 (2020).

2. Quick, J. et al. Multiplex PCR method for MinION and Illumina sequencing of Zika and other virus genomes directly from clinical samples. Nat. Protoc. 12, 1261–1276 (2017).

3. Grubaugh, N. D. et al. An amplicon-based sequencing framework for accurately measuring intrahost virus diversity using PrimalSeq and iVar. Genome Biol. 20, 8 (2019).

4. XSEDE. www.xsede.org.

5. ELIXIR-DE. https://www.denbi.de/elixir-de.

6. ELIXIR. https://elixir-europe.org/.

7. https://nectar.org.au/.

8. Goecks, J., Nekrutenko, A., Taylor, J. & Team, G. Galaxy: a comprehensive approach for supporting accessible, reproducible, and transparent computational research in the life sciences. Genome Biol. 11, R86 (2010).

9. Blankenberg, D., Taylor, J., Nekrutenko, A. & Team, G. Making whole genome multiple alignments usable for biologists. Bioinformatics 27, 2426–2428 (2011).

10. Batut, B. et al. Community-Driven Data Analysis Training for Biology. Cell Syst 6, 752–758.e1 (2018).

11. Grüning, B. et al. Bioconda: sustainable and comprehensive software distribution for the life sciences. Nat. Methods 15, 475–476 (2018).

12. Grüning, B. A. et al. Jupyter and Galaxy: Easing entry barriers into complex data analyses for biomedical researchers. PLoS Comput. Biol. 13, e1005425 (2017).

13. Galactic Introductions. http://www.youtube.com/playlist?list=PLNFLKDpdM3B9UaxWEXgziHXO3k-003FzE.

14. Wilm, A. et al. LoFreq: a sequence-quality aware, ultra-sensitive variant caller for uncovering cell-population heterogeneity from high-throughput sequencing datasets. Nucleic Acids Res. 40, 11189–11201 (2012).

15. Itokawa, K., Sekizuka, T., Hashino, M., Tanaka, R. & Kuroda, M. Disentangling primer interactions improves SARS-CoV-2 genome sequencing by multiplex tiling PCR. PLoS One 15, e0239403 (2020).

16. Lemieux, J. et al. Phylogenetic analysis of SARS-CoV-2 in the Boston area highlights the role of recurrent importation and superspreading events. medRxiv (2020) doi:10.1101/2020.08.23.20178236.

17. du Plessis, L. et al. Establishment and lineage dynamics of the SARS-CoV-2 epidemic in the UK. Science 371, 708–712 (2021).

18. arambaut, garmstrong & isabel. Preliminary genomic characterisation of an emergent SARS-CoV-2 lineage in the UK defined by a novel set of spike mutations. https://virological.org/t/preliminary-genomic-characterisation-of-an-emergent-sars-cov-2-lineage-in-the-uk-defined-by-a-novel-set-of-spike-mutations/563/2 (2020).

19. PANGO lineages. https://cov-lineages.github.io/lineages-website/global_report.html.

20. Greaney, A. J. et al. Comprehensive mapping of mutations to the SARS-CoV-2 receptor-binding domain that affect recognition by polyclonal human serum antibodies. Cold Spring Harbor Laboratory 2020.12.31.425021 (2021) doi:10.1101/2020.12.31.425021.

21. Kosakovsky Pond, S. L. & Frost, S. D. W. Not So Different After All: A Comparison of Methods for Detecting Amino Acid Sites Under Selection. Mol. Biol. Evol. 22, 1208–1222 (2005).

22. Murrell, B. et al. Detecting Individual Sites Subject to Episodic Diversifying Selection. PLoS Genet. 8, e1002764 (2012).

23. Garrison, E. & Marth, G. Haplotype-based variant detection from short-read sequencing. arXiv.org q-bio.GN, (2012).

24. DePristo, M. A. et al. A framework for variation discovery and genotyping using next-generation DNA sequencing data. Nat. Genet. 43, 491–498 (2011).

25. Barrick, J. E. et al. Genome evolution and adaptation in a long-term experiment with Escherichia coli. Nature 461, 1243–1247 (2009).

26. Wei, Z., Wang, W., Hu, P., Lyon, G. J. & Hakonarson, H. SNVer: a statistical tool for variant calling in analysis of pooled or individual next-generation sequencing data. Nucleic Acids Res. 39, e132 (2011).

27. Salk, J. J., Schmitt, M. W. & Loeb, L. A. Enhancing the accuracy of next-generation sequencing for detecting rare and subclonal mutations. Nat. Rev. Genet. 19, 269–285 (2018).

28. Bush, S. J. et al. Genomic diversity affects the accuracy of bacterial SNP calling pipelines. bioRxiv 653774 (2019) doi:10.1101/653774.

29. Yoshimura, D. et al. Evaluation of SNP calling methods for closely related bacterial isolates and a novel high-accuracy pipeline: BactSNP. Microbial Genomics 5, e000261 (2019).

30. Mei, H., Arbeithuber, B., Cremona, M. A., DeGiorgio, M. & Nekrutenko, A. A High-Resolution View of Adaptive Event Dynamics in a Plasmid. Genome Biol. Evol. 11, 3022–3034 (2019).

31. Schmitt, M. W. et al. Detection of ultra-rare mutations by next-generation sequencing. Proc. Natl. Acad. Sci. U. S. A. 109, 14508–14513 (2012).

